# Validation of Solvent Proteome Profiling for Antimalarial Drug Target Deconvolution

**DOI:** 10.1101/2025.08.15.670428

**Authors:** Yijia Ji, Joshua P. Morrow, Christopher A. MacRaild, Haijian Zhang, Carlo Giannangelo, Ralf B. Schittenhelm, Darren J. Creek, Ghizal Siddiqui

## Abstract

Malaria remains a global health threat, with rising drug resistance accelerating the urgent need for new therapeutics. Target elucidation is a critical step in antimalarial drug discovery, enabling a deeper understanding of the molecular mechanisms of action of both existing and novel compounds. This study validates solvent-induced proteome profiling (SPP) as a proteomics-based approach for identifying drug-protein interactions in *Plasmodium falciparum*. SPP detects shifts in protein stability induced by ligand binding, allowing the identification of drug target/s without the need for compound modifications. Here, we successfully generated solvent denaturation curves for the *P. falciparum* proteome, and demonstrated the utility of SPP with five antimalarial compounds: pyrimethamine, atovaquone, cipargamin, MMV1557817 and OSM-S-106. In addition to measuring each compound’s impact across the full denaturation curve, investigating protein levels at individual solvent percentages preserved specific stability changes that would otherwise be masked in pooled analyses performed by integral SPP. This strategy was critical for the identification of the cipargamin target, non-SERCA-type Ca^2+^-transporting P-ATPase (*Pf*ATP4). Notably, we propose live-cell treatment SPP as a novel approach, demonstrating its ability to identify the validated target of pyrimethamine, bifunctional dihydrofolate reductase-thymidylate synthase (*Pf*DHFR), with high specificity. We also introduced the novel one-pot mixed-drug SPP which enables the evaluation of multiple drugs within a single lysate and experimental setup. This alternative method simplifies the experimental workflow and includes positive controls to affirm the performance of the experiment. Overall, this study demonstrates that SPP can be successfully applied in both lysates and live cells to elucidate drug targets in *P. falciparum,* as well as providing additional information regarding the mechanisms of drug action, offering insights for the optimisation of existing antimalarials and the development of novel therapies.

**Graphical Abstract:** 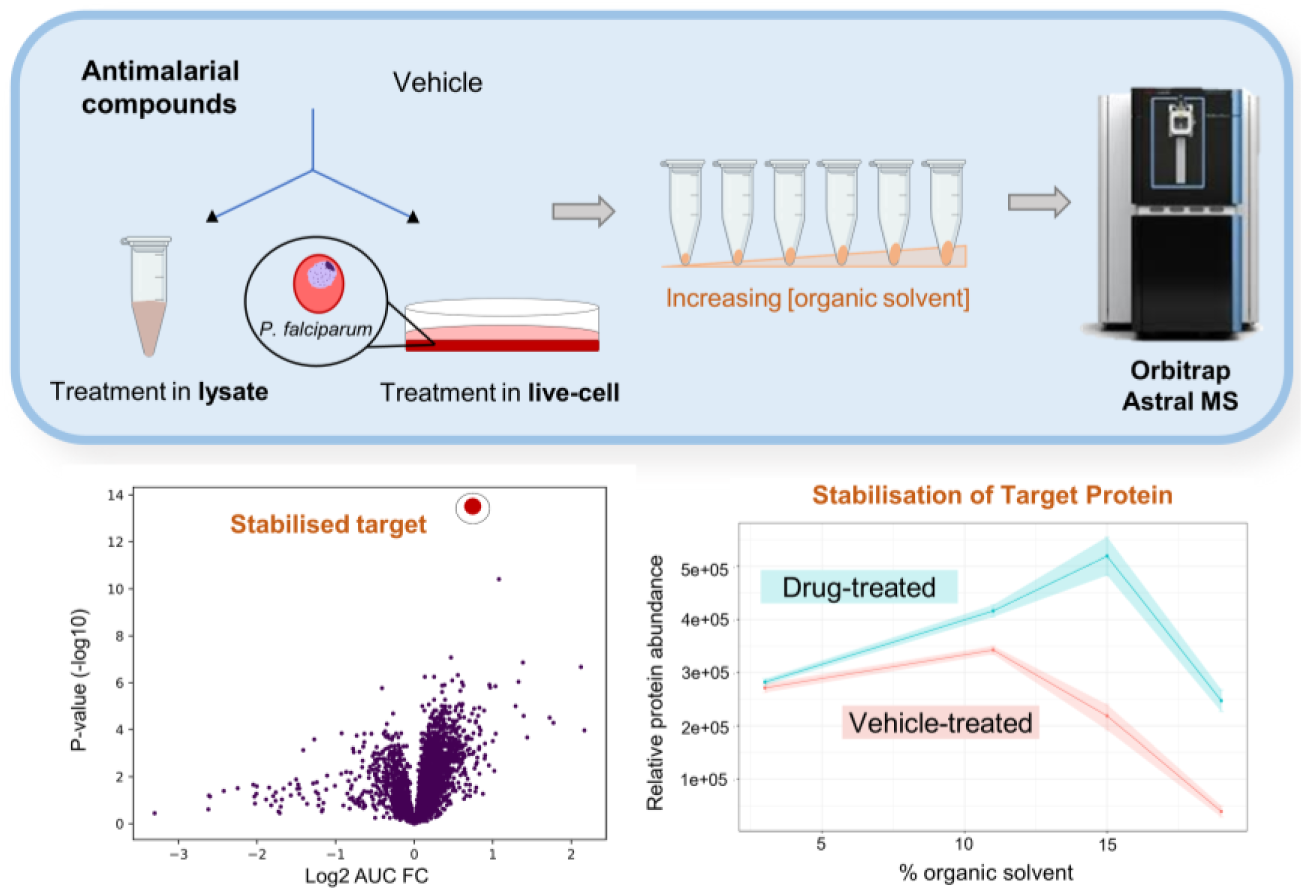

## 1. Introduction

Exploring the interactions between proteins and ligands for drug target identification (target ID) is a crucial step in the drug discovery and development pipeline. Phenotypic screening is a common drug discovery method that prioritises observable biological activity without direct evaluation of the target responsible for the activity (Challis et al., 2022). In the past decade, over 50% of clinical trials failed due to lack of drug efficacy (Fogel, 2018). This could be partially attributed to poor target validation and insufficient understanding of a drug’s mechanism of action (MOA) (Trowbridge et al., 2022). Malaria exemplifies this problem, where many promising drug candidates, including the current first-line artemisinins (ART), have emerged based on their phenotypic activity without a known target. This lack of target knowledge not only hampers drug development, but also restricts the capacity to anticipate resistance (Challis et al., 2022). Without target knowledge, rational design of combination therapies to prevent or delay resistance becomes nearly impossible. Consequently, resistance has emerged against many antimalarials in the field (Gregson and Plowe, 2005; Wellems and Plowe, 2001; Wongsrichanalai et al., 2002), including artemisinin-based combination therapies (ACTs) (Amaratunga et al., 2016; Balikagala et al., 2021; Mairet-Khedim et al., 2021; Phyo et al., 2016; Rosenthal et al., 2024; Tumwebaze et al., 2022). As a repercussion, malaria cases reached 263 million in 2023 - a significant increase from 200 million cases in 2015 - alongside a rise in deaths from 400, 000 to approximately 600, 000 in the same period (WHO, 2014; WHO, 2024). This stalling in progress against malaria underscores the urgent need for advanced methods to explore protein targets of both existing and novel antimalarials. This can improve clinical translation, achieved through effective target deconvolution and optimisation of candidates against the identified target(s) (Umumararungu et al., 2023).

Target deconvolution requires powerful tools to probe target engagement within a whole proteome. Conventional proteomics-based target ID methods, such as protein pull-down assays and affinity chromatography, rely on a chemically modified ligand to isolate interacting partners from complex mixtures (Challis et al., 2022). Though effective, these strategies require chemical modification of the compound of interest, which can affect its activity or membrane permeability (Klein et al., 2020), moving away from native drug-protein interactions (Ha et al., 2021). Chemical modification of novel drug candidates also introduces challenges in chemical synthesis, purification and stability, requiring intricate experimental design and technical expertise, and may not be feasible for all compounds (Liu et al., 2022). Additionally, these methods also identify many non-specifically binding proteins (Challis et al., 2022).

Recent advances in mass spectrometry (MS) have provided pharmaceutical researchers with enhanced capability in quantitative proteomics using label-free methods to investigate proteome-wide drug-protein interactions (Challis et al., 2022). Techniques such as the cellular thermal shift assay (CETSA) (Jafari et al., 2014) and thermal proteome profiling (TPP) (Savitski et al., 2014) have been widely used to monitor drug-induced protein stabilisation upon heat denaturation. Another related approach, stability of proteins from rates of oxidation (SPROX), applies redox-based chemical denaturation to assess shifts in protein stability following drug treatment (Strickland et al., 2013). Limited proteolysis coupled with mass spectrometry (LiP-MS) detects structural alterations in proteins upon ligand binding, but can only be used for proteins with sufficiently exposed regions for proteolytic digestion. More critically, the drug-ligand interaction site itself must be surface-accessible for structural changes to be detected (Reber and Gstaiger, 2023; Schopper et al., 2017; Tsai et al., 2002). Solvent-induced proteome profiling (SPP) is a newer label-free technique that builds on the same principles of detecting protein stability shifts upon drug binding, in this case following exposure to a denaturing solvent (Van Vranken et al., 2021; Zhang et al., 2020). In SPP, the process of protein unfolding by organic solvents is influenced by drugtarget engagement, which alters the target’s susceptibility to destabilisation/stabilisation. Thus, by analysing the differences in denaturation curves between ligand-treated and vehicle control proteomes, SPP allows protein targets to be identified from complex mixtures. SPP offers an attractive alternative to TPP, particularly for proteins that are less responsive to heat denaturation (Van Vranken et al., 2021). Furthermore, it captures a broader range of protein-protein interactions (PPIs) than LiP-MS.

Typically, whole-proteome stability-based assays are interrogated in cell lysates (Bravo et al., 2024; Van Vranken et al., 2021; Zhang et al., 2020), although this is a compromise as it takes proteins out of their native environment and may not include membrane proteins that cannot be readily extracted in native conditions. Therefore, the study of drug-protein interactions in live cells holds great value. This is critical for some antimalarials, particularly those requiring modification or activation to exert their effects. Certain antimalarials, such as ARTs, depend on parasite-specific cellular processes for their activation (Siddiqui and Giannangelo et al., 2022), highlighting the importance of studying live-cell systems, where the dynamic interaction of metabolic and enzymatic processes is essential for drug activity. In addition, live-cell conditions enable both the identification of direct drug targets and indirect downstream pathways affected as a result of drug exposure, providing deeper insights into the MOA. TPP/CETSA have been adapted for live-cell analysis and have demonstrated their utility in mapping drug-target interactions within intact human cells and parasites (Dziekan et al., 2020; Savitski et al., 2014). Recently, Bravo et al. (2024) has applied SPP to study antimalarial compounds, demonstrating its utility in identifying drug targets in parasite lysates. However, no studies to date have tested SPP in live *P. falciparum* parasites for antimalarial compounds; thus, in this study we have utilised live-cell treatment SPP to study drug-target interactions within intact cells and investigated interactions that depend on specific biological environments. Previous studies in cell lysates have typically adopted integral SPP (iSPP, also known as solvent-proteome integral solubility alteration), which pools soluble protein fractions across solvent concentrations to investigate differences in total abundance between control and drug-treated conditions (Bravo et al., 2024; Van Vranken et al., 2021). Although iSPP simplifies analysis and reduces instrument time, it risks overlooking subtle or solvent-specific stabilisation patterns and may miss weak or transient ligand-protein interactions (Zhang et al., 2023). Moreover, without examining full destabilisation profiles, false positives can arise (Bravo et al., 2024). To address these limitations, we maintained separate solvent conditions and analysed stability changes at individual solvent concentrations. To balance this with experimental efficiency, we also developed a one-pot, mixed-drug lysate SPP strategy, which enables simultaneous detection of multiple ligand-target interactions in a single lysate while retaining the destabilisation curve.

Accurate target engagement studies with precise label-free quantitative data are needed in the field of antimalarial drug target deconvolution. In this study, we applied the latest high-resolution LC/MS-MS methodology for SPP-based drug target deconvolution in *P. falciparum* lysates and live cultures, incorporating optimised solvent concentrations and a one-pot, mixed-drug strategy to enhance target detection. We validated SPP’s robustness in identifying known and unknown drug targets of pyrimethamine (PYR), atovaquone (ATV), cipargamin (CIP), MMV1557817 (Vinh et al., 2019) and OSM-S-106 (Gamo et al., 2010). By establishing this approach, we aim to contribute to drug optimisation by identifying key targets of novel compounds, which can then guide the design of new antimalarials with improved efficacy and reduced risk of resistance.

## 2. Materials and methods

### 2.1 Chemicals and reagents

Pyrimethamine and atovaquone were purchased from SelleckChem and AK Scientific, respectively. Cipargamin and MMV1557817 were purchased from MedChemExpress. Asn-OSM-S-106 was kindly provided by Prof Matthew Todd, University College London. Unless otherwise stated, all reagents were purchased from Merck Life Science Pty Ltd.

### 2.2 *P. falciparum* cell culture and preparation

Asexual *P. falciparum* (3D7 strain) was cultured as previously described (Trager and Jensen, 1976), and parasites were tightly synchronised by double sorbitol lysis to trophozoite stage (34-38 h post invasion). For lysate SPP, parasites were released from the host red blood cells (RBCs) using hypotonic lysis buffer (150 mM ammonium chloride, 10 mM potassium bicarbonate, 1 mM ethylenediaminetetraacetic acid), centrifuged and collected before being frozen at -80 °C for future use. For live-cell treatment SPP, parasites were magnet-harvested as previously described (Ribaut et al., 2008) and subjected to drug or vehicle treatments.

### 2.3 Preparation of drug-treated parasite cell lysate

Parasite pellets collected from hypotonic lysis (synchronised to the trophozoite stage (34-38 h post invasion)) were resuspended in 1 mL of lysis buffer (containing 1× phosphate-buffered saline (PBS), 1.5 mM magnesium chloride (MgCl₂), 150 mM potassium chloride (KCl), 0.4% Nonidet P-40 (NP-40), 250 U/mL Benzonase® Nuclease, protease inhibitor [Pierce Protease Inhibitor Mini Tablets] and phosphatase inhibitor [Roche PhosSTOP^TM^ Tablets]) with shaking at 4 °C for 20 min before centrifugation at 20, 000 × *g* for 10 min at 4 °C to obtain the soluble proteome. Protein lysate concentration was measured using Pierce™ Dilution-Free™ Rapid Gold BCA Protein Assay Kit (Thermo Fisher Scientific) and diluted to a concentration of 2 mg·mL^-1^. Supernatant was then divided into equal aliquots (*n* = 2-3) for vehicle (dimethyl sulfoxide (DMSO)) and compound treatment (1 µM) and was incubated at room temperature for 15 min. Treated lysate was then used for subsequent SPP experiments.

### 2.4 Preparation of drug-treated live parasite

Magnet harvested parasites (80-90% infected red blood cells (iRBCs), synchronised to the trophozoite stage (34-38 h post invasion)) were incubated in media containing the drug (1 µM of PYR for 30 min) or vehicle control. Following drug incubation, cells were washed twice with PBS and lysed in ice-cold lysis buffer (1× PBS, 1.5 mM MgCl₂, 150 mM KCl, 0.8% NP-40, 250 U/mL Benzonase® Nuclease, protease inhibitor [Pierce Protease Inhibitor Mini Tablets] and phosphatase inhibitor [Roche PhosSTOP^TM^ Tablets]) for 30 min at 4 °C with agitation (2000 rpm). Haemoglobin was depleted by incubating lysates with 500 µL Talon® Metal Affinity Resin (Takara) pre-washed with PBS at room temperature for 5 min, followed by centrifugation (500 × g, 30 s). This step was repeated 3-4 times until supernatants were clear. Lysate protein concentration was measured using Pierce™ Dilution-Free™ Rapid Gold BCA Protein Assay Kit (Thermo Fisher Scientific) and diluted to a concentration of 2 mg·mL^-1^ if needed. The supernatant containing the proteome was used for subsequent SPP and analysis.

### 2.5 Solvent-induced proteome profiling

Solvent-induced protein precipitation and proteome profiling was done as previously described (Van Vranken et al., 2021; Zhang et al., 2020). An equal volume of the lysate from drug/vehicle treated conditions (from live cell or cell lysates) was added to tubes containing additional lysis buffer and a mixture of acetone/ethanol/acetic acid (50:50:0.1 v/v/v, AEA). For initial curve generation in solvent-induced proteome precipitation, %AEA was adjusted to 0, 1, 3, 5, 9, 11, 13, 15, 17, 19, 21, 23, 25 (*n* = 2-4). For lysate SPP with PYR and ATV, as well as live-cell treatment SPP with PYR, solvent percentages of 3, 11, 15, 19 were applied. For lysate SPP with CIP, solvent percentages of 3, 9, 11, 13, 15, 19 were applied. For lysate one-pot SPP, solvent percentages of 5, 10, 13, 15, 23 were applied (*n* = 2-3). Upon the addition of AEA, the samples were incubated at 37 °C with vigorous shaking for 20 min and then centrifuged at 4 °C (21, 000 × *g*, 15 min). The supernatant containing the soluble protein fraction was taken (85 µL) for reduction (10 mM tris(2-carboxyethyl)phosphine hydrochloride (Pierce^TM^)) and alkylation (40 mM 2-chloroacetamide) for 15 min at 55 °C.

The samples were then subjected to ProtiFi S-trap^™^ mini clean-up steps according to the manufacturer’s protocol with trypsinisation (added at a ratio of 1:50 enzyme/protein w/w) and desalted using in-house generated C18 StageTips (ProtiFi; Rappsilber et al., 2003). Samples were then dried, and resuspended in 12 μL of 2% acetonitrile (ACN) and 0.1% formic acid (FA), spiked with indexed retention time (iRT) peptides for LC-MS/MS analysis.

### 2.6 LC-MS/MS analysis with Orbitrap Astral using LFQ-DIA

Samples were analysed on a Vanquish Neo HPLC coupled to an Orbitrap Astral mass spectrometer (Thermo Fisher Scientific) equipped with NanoSpray Flex source (Thermo Fisher Scientific), using a 50 cm μPAC^™^ HPLC column (2.5 μm, 5 × 28 μm^2^, C18, 100-200 Å, Thermo Fisher Scientific) or a 25 cm Aurora Ultimate™ XT UHPLC column (1.7 μm, 75 µm, C18, 120 Å, IonOpticks). The CIP single-drug experiment was performed using the 25 cm Aurora Ultimate™ XT UHPLC column, while the live-cell treatment, one-pot, and single-drug experiments with ATV and PYR were conducted using the 50 cm μPAC™ HPLC column. Mobile phase A and B were 0.1% FA in water (Thermo Fisher Scientific, Optima LC-MS grade) and 0.1% FA/80% ACN (Thermo Fisher Scientific, Optima LC-MS grade), respectively. The column was heated to 50 °C, with the flow rate initially set to fast load (dependent on the column pressure limit of 700 nL/min and 450 bar) to minimise delay time. At the start of the active gradient, the flow rate was turned to 350 nL/min. The initial conditions of 4% B were ramped up to 30% from 0 to 45 min, followed by an increase to 45% over 15 min. The column was then washed for 5 min at 97.5% B with a flow rate of 700 nL/min at the end of the active gradient, followed by fast equilibration on the Vanquish Neo LC, with a maximum pressure limit of 450 bar. For data independent acquisition (DIA) experiments on the Orbitrap Astral MS, MS1 spectra were collected in the Orbitrap at a resolving power of 240, 000 over an *m/z* of 380-980. The MS1 normalised AGC target was set to 500% with a maximum injection time of 5 ms. DIA MS2 scans were acquired using the Astral analyser, covering the 380-980 *m/z* range, with a normalised AGC target of 800%, a maximum injection time of 3 ms and an HCD collision energy of 25%, with a default charge state of +2. Window placement optimisation was enabled, with isolation widths of 2 Th and active gradient length of 60 min.

### 2.7 Analysis of SPP LC-MS/MS data

The proteomics data was processed using Spectronaut® Pulsar (v19.0, Biognosys, Switzerland) (version 19.2.240905.62635) referencing an in-house *P. falciparum* protein spectral library as per default settings (Siddiqui et al., 2022). The data files exported from Spectronaut were analysed and visualised in DIA-Analyst (https://analyst-suites.org/apps/dia-analyst/), which is currently in the process of undergoing publication elsewhere. The data was further examined using CurveLine (https://analyst-suites.org/apps/curveline/), which is our publicly accessible web application specifically developed to assess individual protein abundances in SPP data across various conditions. Detailed documentation and description of the CurveLine tool are available on our GitHub repository (https://github.com/MonashProteomics/CurveLine). Both applications automatically processed the data from Spectronaut®, identified proteins and quantified protein abundances. Briefly, proteins lacking quantitative values (reverse sequences, potential contaminant sequences, proteins identified solely by site) were filtered from the dataset to ensure robustness of subsequent analyses. Following pre-processing, all intensity values were log_2_-transformed and grouped according to solvent and drug conditions. Protein-wise linear models incorporating empirical Bayes statistics were applied to perform differential expression analyses. Utilising the Bioconductor package limma, significantly stabilised proteins in each pairwise comparison were identified using a threshold of *p* ≤ 0.05, controlled by the Benjamini-Hochberg (BH) method (Monash Proteomics and Metabolomics Facility, 2024). Additionally, proteins were considered to show significant stabilisation only if they met both the *p*-value and log_2_ fold change (log_2_FC) thresholds between control and experimental groups. Area under the curve (AUC) volcano plots were generated by first computing AUC for each SPP profile, which the sum of the mean protein abundance was approximated at each AEA percentage. Uncertainties on each AUC were estimated by calculating the relative standard deviation (RSD) at each AEA percentage and taking the average RSD across each profile as the effective RSD for the AUC. *p*-value was adjusted using the BH method.

## 3. Results

### 3.1 Experimental overview

To deconvolute the targets of known antimalarial compounds, we applied SPP using the Orbitrap Astral MS with the label-free quantification approach and data-independent acquisition (LFQ-DIA). This approach was used to validate and potentially identify additional drug targets for a range of known antimalarials in both live parasites and lysates. Our process also streamlined data analysis using DIA-Analyst, a web application for differential expression analysis and visualisation of LFQ proteomic datasets preprocessed from Spectronaut®. Additionally, we created a publicly accessible web application entitled ‘CurveLine’ to examine trends in individual protein abundance across solvent percentages and conditions.

### 3.2 Solvent-induced proteome precipitation (SIP) of *P. falciparum* parasite lysates

To assess the application of SPP in *P. falciparum*, 3D7 parasite lysates (*n* = 2-4) were treated with increasing concentrations of AEA (0-25%) and subjected to LFQ-DIA quantitative proteomics to investigate the denaturation patterns of the *P. falciparum* iRBC proteome. A total of 4, 674 proteins were detected, including 3, 839 *P. falciparum* proteins. Over 98% of proteins were identified with 2 or more peptides, demonstrating robust proteomic coverage and the detection of an unprecedented number of *P. falciparum* proteins across samples (Figure 1A).

**Figure 1.**
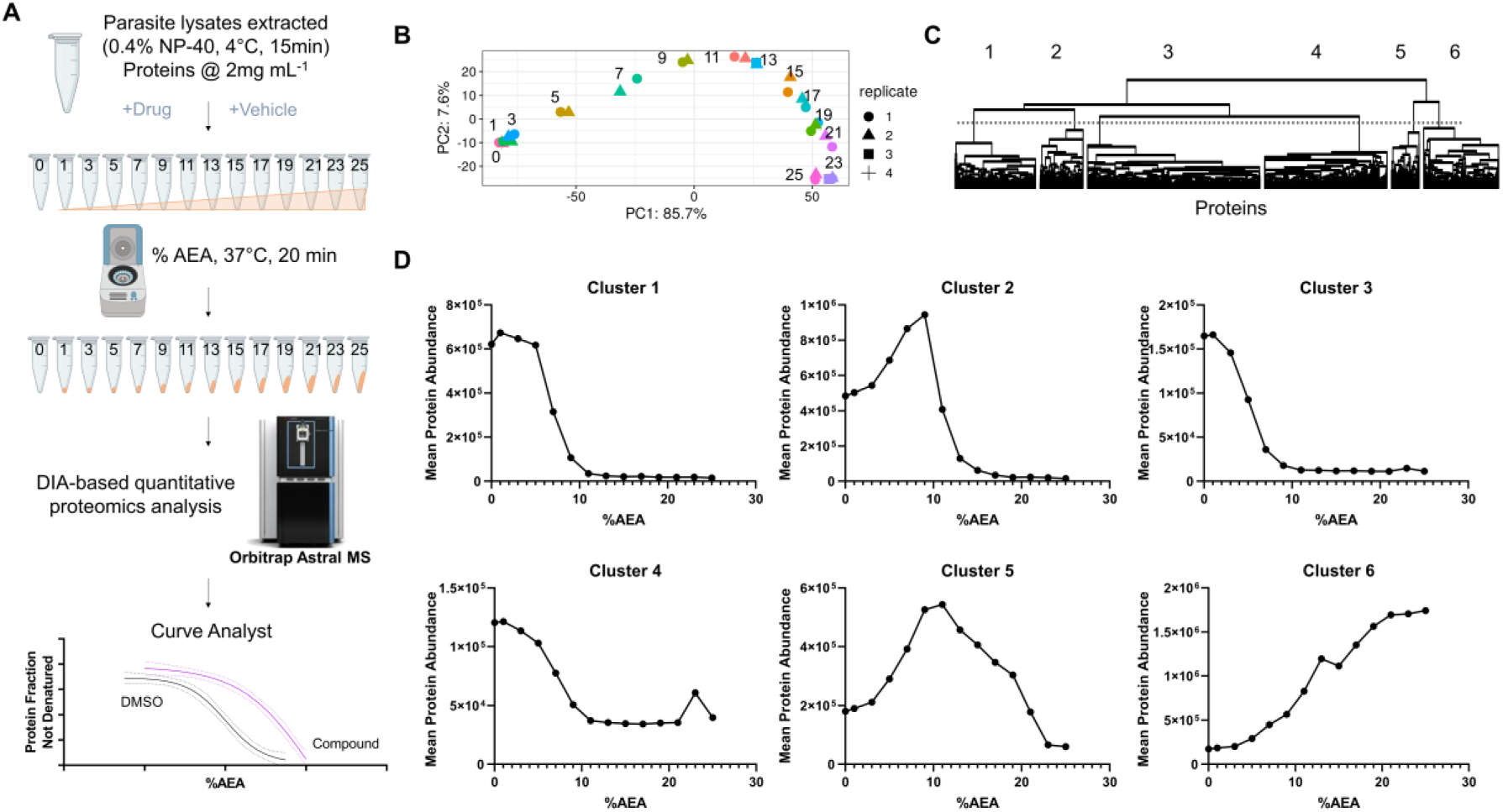
Solvent-induced proteome precipitation of *P. falciparum* lysate. (A) Schematic overview of the SPP workflow using parasite lysate. Briefly, parasite lysates were extracted and protein concentration was adjusted accordingly. Lysates were treated with vehicle or drugs, and exposed to a mixture of acetone/ethanol/acetic acid (50:50:0.1 v/v/v, AEA) at 0, 1, 3, 5, 7, 9, 11, 13, 15, 17, 19, 21, 23 and 25%. Soluble proteins were recovered by centrifugation, purified, and subjected to downstream proteomic analysis with the Orbitrap Astral MS. MS data were processed and analysed as described to observe shifts in protein abundance curves. (B) Principal component analysis of the global proteome revealed clear separation of samples along PC1, corresponding to altered protein levels with increasing %AEA. (C) Hierarchical clustering analysis of protein abundance with increasing %AEA (0-25%) reveal six distinct protein clusters. The dendrogram groups proteins based on similar abundance changes in response to AEA. Figure 1B and Figure 1D were generated using DIA analyst (https://analyst-suites.org/apps/dia-analyst/). (D) Curve plots showing the mean abundance of all proteins within each cluster in response to AEA at 0, 1, 3, 5, 7, 9, 11, 13, 15, 17, 19, 21, 23 and 25%. Clusters 1-5 show decreasing abundance with increasing %AEA, while cluster 6 shows the opposite pattern, with protein abundance increasing as %AEA increases. The plots were created using GraphPad Prism 10.

Principal component analysis (PCA) revealed clear separation of sample groups based on AEA concentration. The first principal component (PC1), accounting for 85.7% of the variance, showed clear separation between solvent percentages (Figure 1B). Additionally, hierarchal clustering analysis further confirmed these trends, showing systematic changes in soluble protein abundance with increasing solvent percentages (Figure 1C, Figure 1D). Interestingly, while most proteins decreased in abundance as solvent percentage increased, as expected due to protein denaturation and precipitation, proteins in cluster 6 were detected with greater abundance in the higher %AEA samples (Figure 1D). Nevertheless, these patterns suggest that AEA effectively alters protein abundance in a concentration-dependent manner, providing a basis for identifying proteins stabilised or destabilised following drug treatment. The SPP workflow presented here serves as a foundation for subsequent drug-target engagement studies under live-cell treated and lysate conditions.

### 3.3 Drug-induced protein stabilisation in *P. falciparum* parasite lysates

To evaluate the ability of the SPP approach to identify the known targets and perhaps novel targets of current antimalarials: PYR, ATV and CIP in *P. falciparum* lysates, 3D7 parasite protein lysates were subjected to increasing AEA concentrations (3%, 9%, 11%, 13%, 15%, 19%) in the presence of either DMSO or the respective drugs (1 μM for 15 min). Previous studies have demonstrated that some proteins exhibit differential stabilisation at specific solvent percentages, rather than uniform stabilisation across all conditions (Van Vranken et al., 2021; Zhang et al., 2020). Based on this, stabilisation profiles were analysed based on the AUC from all solvent percentages, and at each solvent percentage individually.

Firstly, PYR was selected due to its well-characterised MOA and established target, bifunctional dihydrofolate reductase-thymidylate synthase (*Pf*DHFR, PF3D7_0417200), making it an ideal validation candidate (Peterson et al., 1988). Hierarchal clustering and heatmap analysis (Figure 2A) revealed distinct patterns of protein abundances as solvent percentage increased and clustering within solvent percentage groups revealed impacts of PYR on protein stabilisation or destabilisation. Volcano plot analysis of AUC of all percentages (vehicle vs drug) identified *Pf*DHFR to be significantly stabilised following incubation with PYR in parasite lysates (*p.adj* = 2.27e-14, log_2_FC = 0.74) (Figure 2B). In Figure 2C, the curve plot showed a marked shift in protein abundance with PYR treatment across increasing %AEA with the most substantial stabilisation observed at 15% AEA. Importantly, *Pf*DHFR was the most significantly stabilised protein (*p.adj* = 2.27e-14), with a considerably lower *p* value than the next most significant proteins PF3D7_1110600 (*p.adj* = 3.86e-11), and PF3D7_1142800 (*p.adj* = 8.43e-08), demonstrating clear identification of the drug target from the parasite proteome.

**Figure 2.**
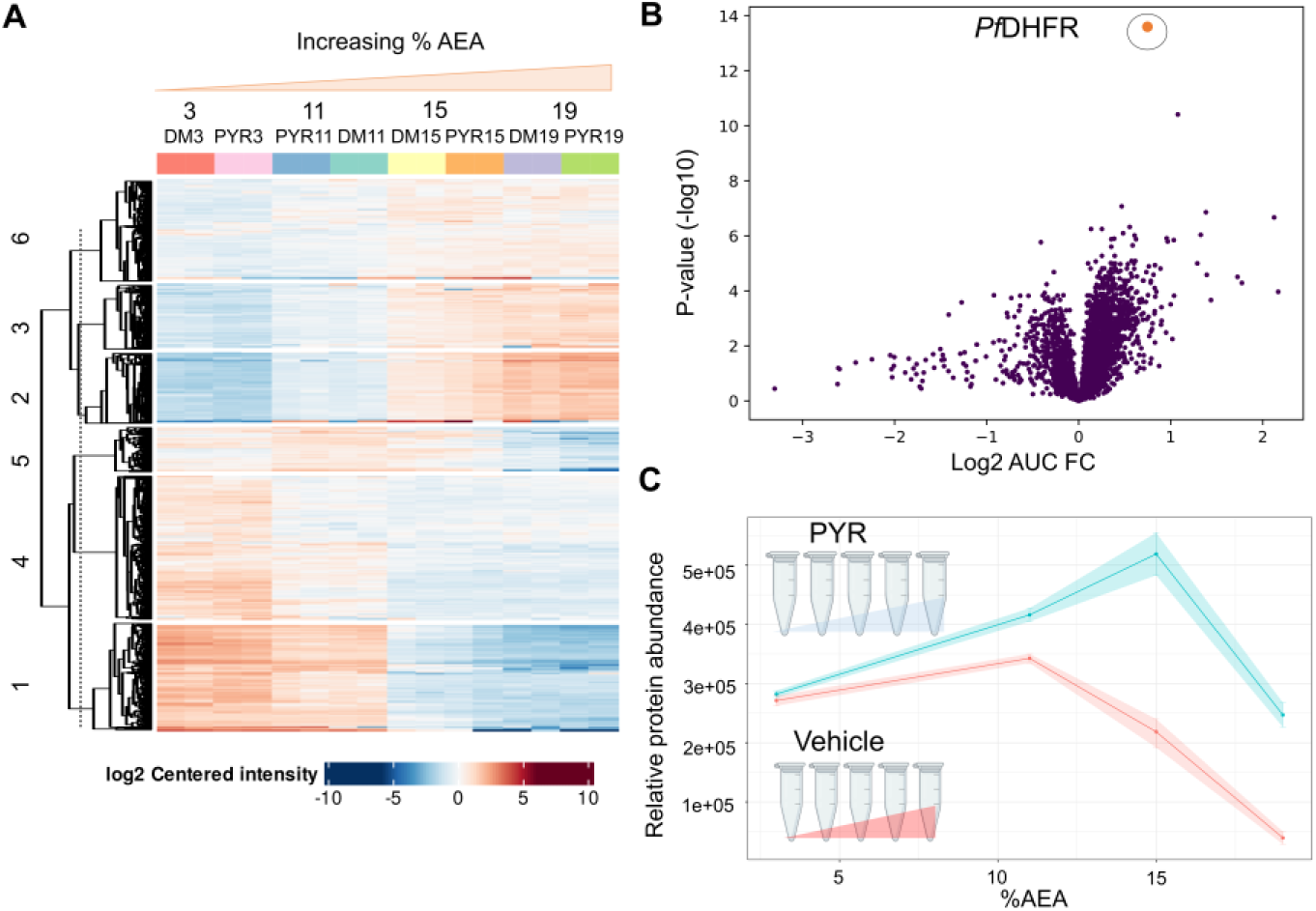
SPP target validation of PYR in parasite lysates. (A) Hierarchical clustering heatmap of parasite lysates treated with increasing concentrations of AEA (3%, 11%, 15%, 19%) in the presence of DMSO (vehicle control) or PYR (1 µM). Each colour block represents an experimental condition. (B) Volcano plot generated from AUC analysis comparing vehicle and PYR-treated samples, highlighting significantly perturbed proteins. *Pf*DHFR (PF3D7_0417200), the protein target of PYR, was identified with a log₂FC of 0.74 and *p.adj* of 2.27e-14. (C) Destabilisation curves of PfDHFR with increasing %AEA, after PYR and DMSO treatments. Protein abundances were plotted against AEA concentrations at 3%, 11%, 15%, 19%. The shaded areas around each curve represent the range between the 2 replicates at each AEA concentration.

### 3.4 Validating cytochrome bc_1_ Complex III as an atovaquone target in *P. falciparum* lysates

Whilst PYR provided an ideal proof of concept for our SPP approach, we also wanted to validate our technique with another known antimalarial, ATV, which targets the cytochrome bc_1_ Complex III of the parasite mitochondria (Fry and Pudney, 1992) and has well-established efficacy (Srivastava et al., 1997). Differential protein destabilisation was observed comparing ATV and DMSO conditions with increasing %AEA (Figure 3A). The most significantly stabilised proteins following incubation with ATV in parasite lysates are highlighted in the AUC volcano plot (vehicle vs drug) in Figure 3C, which included five cytochrome bc_1_ Complex III canonical subunits (cytochrome bc_1_ complex putative subunit 7 (QCR7, PF3D7_1012300, *p.adj* = 1.01e-10, log_2_FC = 2.67); putative subunit 9 (QCR9, PF3D7_0622600, *p.adj* = 2.26e-18, log_2_FC = 1.80); Rieske protein (PF3D7_1439400, *p.adj* = 2.72e-18, log_2_FC = 1.16); cytochrome B (CYTB, PF3D7_MIT02300, *p.adj* = 6.75e-12, log_2_FC = 2.96) and cytochrome c1 (CYTC1, PF3D7_1462700, *p.adj* = 8.33E-13, log_2_FC = 1.77)), as well as four additional proteins (respiratory chain complex 3 associated protein 1 (C3AP1, PF3D7_0722700, *p.adj* = 6.53e-17, log_2_FC = 1.71); respiratory chain complex 3 associated protein 2 (C3AP2, PF3D7_1326000, *p.adj* = 2.72e-18, log_2_FC = 1.70); mitochondrial-processing peptidase subunit alpha (MPPα, PF3D7_0523100, *p.adj* = 2.35e-15, log_2_FC = 1.31), and mitochondrial-processing peptidase subunit beta (MPPβ, PF3D7_0933600, *p.adj* = 9.15e-15, log_2_FC = 1.43)). All nine proteins make up the mitochondrial Complex III in *P. falciparum* as defined by Evers et al. (2021) (Figure 3B), confirming that our SPP technique can be successfully utilised for an antimalarial that targets a protein complex. Further, the stabilisation of the additional subunits, C3AP1, C3AP2, MPPα and MPPβ, confirms their physical association within Complex III, further validating the composition of parasite proteins that make up the cytochrome bc_1_ Complex III.

**Figure 3.**
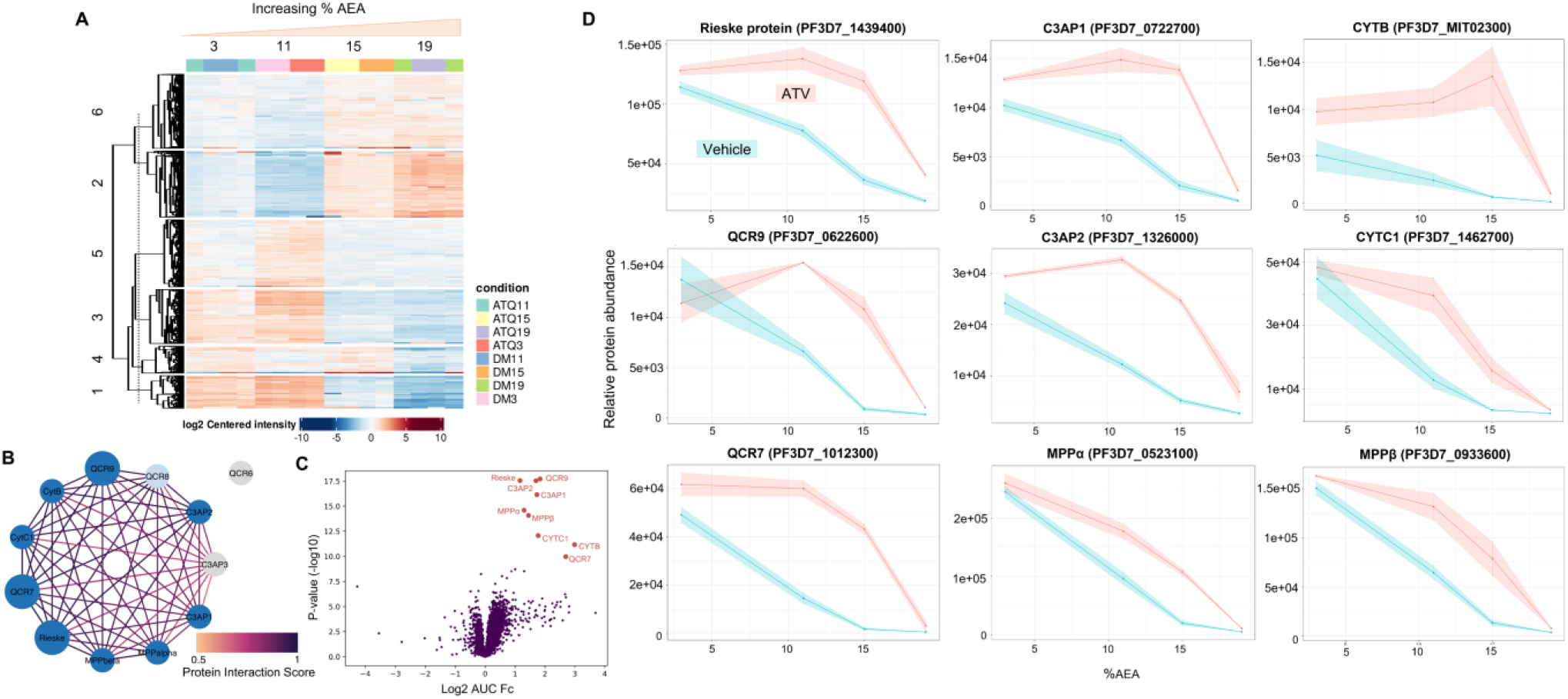
SPP validates ATV protein targets in parasite lysates. (A) Hierarchical clustering heatmap of parasite lysates treated with increasing concentrations of AEA (3%, 11%, 15%, 19%) in the presence of DMSO (vehicle control) or ATV (1 µM). (B) Proposed complexome schematic of parasite cytochrome bc_1_ Complex III, consisting of the cytochrome bc_1_ complex Rieske protein (PF3D7_1439400), cytochrome B (CYTB, PF3D7_MIT02300), cytochrome c1 (CYTC1, PF3D7_1462700), cytochrome bc_1_ complex putative subunit 7 (QCR7, PF3D7_1012300), cytochrome bc_1_ complex putative subunit 9 (QCR9, PF3D7_0622600), respiratory chain complex 3 associated protein 1 (C3AP1, PF3D7_0722700), respiratory chain complex 3 associated protein 2 (C3AP2, PF3D7_1326000), respiratory chain complex 3 associated protein 3 (C3AP3, PF3D7_0817800), mitochondrial-processing peptidase subunit alpha (MPPα, PF3D7_0523100), mitochondrial-processing peptidase subunit beta (MPPβ, PF3D7_0933600), cytochrome bc_1_ complex putative subunit 6 (QCR6, PF3D7_1426900), cytochrome bc_1_ putative complex subunit 8 (QCR8, PF3D7_0306000). Protein interaction scores derived from Evers et al. (2021). Dark blue circles indicate stabilised proteins; light blue circles indicate proteins detected but without significant stability change following ATV exposure; grey circles indicate proteins not detected in our iRBC SPP data. (C) AUC volcano plot of vehicle vs drug with consistently identified significant protein targets (red dots), including the Rieske protein (*p.adj* = 2.72e-18, log_2_FC = 1.16), CYTB (*p.adj* = 6.75e-12, log_2_FC = 2.96), CYTC1 (*p.adj* = 8.33e-13, log_2_FC = 1.77), QCR7 (*p.adj* = 1.01e-10, log_2_FC = 2.67), QCR9 (*p.adj* = 2.26e-18, log_2_FC = 1.80), C3AP1 (*p.adj* = 6.53e-17, log_2_FC = 1.71), C3AP2 (*p.adj* = 2.72e-18, log_2_FC = 1.70), MPPα (*p.adj* = 2.35e-15, log_2_FC = 1.31), MPPβ (*p.adj* = 9.15e-15, log_2_FC = 1.43). Cut-off of the volcano plot is log₂FC > 1 and *p.adj* < 1e-09. (D) Destabilisation curves of Rieske protein, CYTB, CYTC1, QCR7, QCR9, C3AP1, C3AP2, MPPα and MPPβ with increasing %AEA, after ATV and DMSO treatments. Protein abundances were plotted against AEA concentrations at 3%, 11%, 15%, 19%. The shaded areas around each curve represent the range between the 2 replicates at each AEA concentration. Destabilisation curve of QCR8 is included in Figure S2.

### 3.5 Validation of *Pf*ATP4 as a cipargamin target using solvent-specific analysis

In addition to PYR and ATV, we included CIP, an antimalarial that has completed its phase II clinical trials (Schmitt et al., 2021; White et al., 2014), to demonstrate the applicability of our SPP approach to membrane-associated targets. CIP inhibits the parasite plasma membrane-localised non-SERCA-type Ca^2+^-transporting P-ATPase (*Pf*ATP4, PF3D7_1211900) (Spillman and Kirk, 2015), a sodium efflux pump that maintains ionic homeostasis in the parasite (Krishna et al., 2001). Despite the hierarchical clustering heatmap showing proteome-wide trends similar to previously examined antimalarials (Figure 4A), AUC analysis with BH correction yielded no significantly stabilised proteins, including CIP’s known target *Pf*ATP4 (*p.adj* = 0.49, log₂FC = 0.08) (Rottmann et al., 2010). This prompted a closer analysis of target stabilisation at individual %AEA. At 15% AEA, the volcano plot (vehicle vs drug) (Figure 4C) revealed several stabilised proteins, including *Pf*ATP4 that shows significant stabilisation following incubation with CIP in parasite lysates (*p.adj* = 1.6e-05, log₂FC = 0.92). The perturbation of *Pf*ATP4 by CIP was further supported by the destabilisation curve (Figure 4B), where CIP-treated samples showed a significant abundance shift only at 15% AEA compared to DMSO control. This is also illustrated by the bar plots of *Pf*ATP4 abundance in CIP-treated samples at 15% AEA compared to AUC (Figure 4D). This demonstrates that the pooled solvent percentages approach may not be appropriate for all drugs and that analysing individual solvent specific changes across the target stabilisation curves is critical. Two other proteins, merozoite surface protein 1 (MSP1, PF3D7_0930300) and putative rhoptry protein (ROP, PF3D7_0721400) were consistently stabilised across multiple AEA percentages and further studies are required to determine whether these are genuine secondary binding partners of CIP (Figure S2).

**Figure 4.**
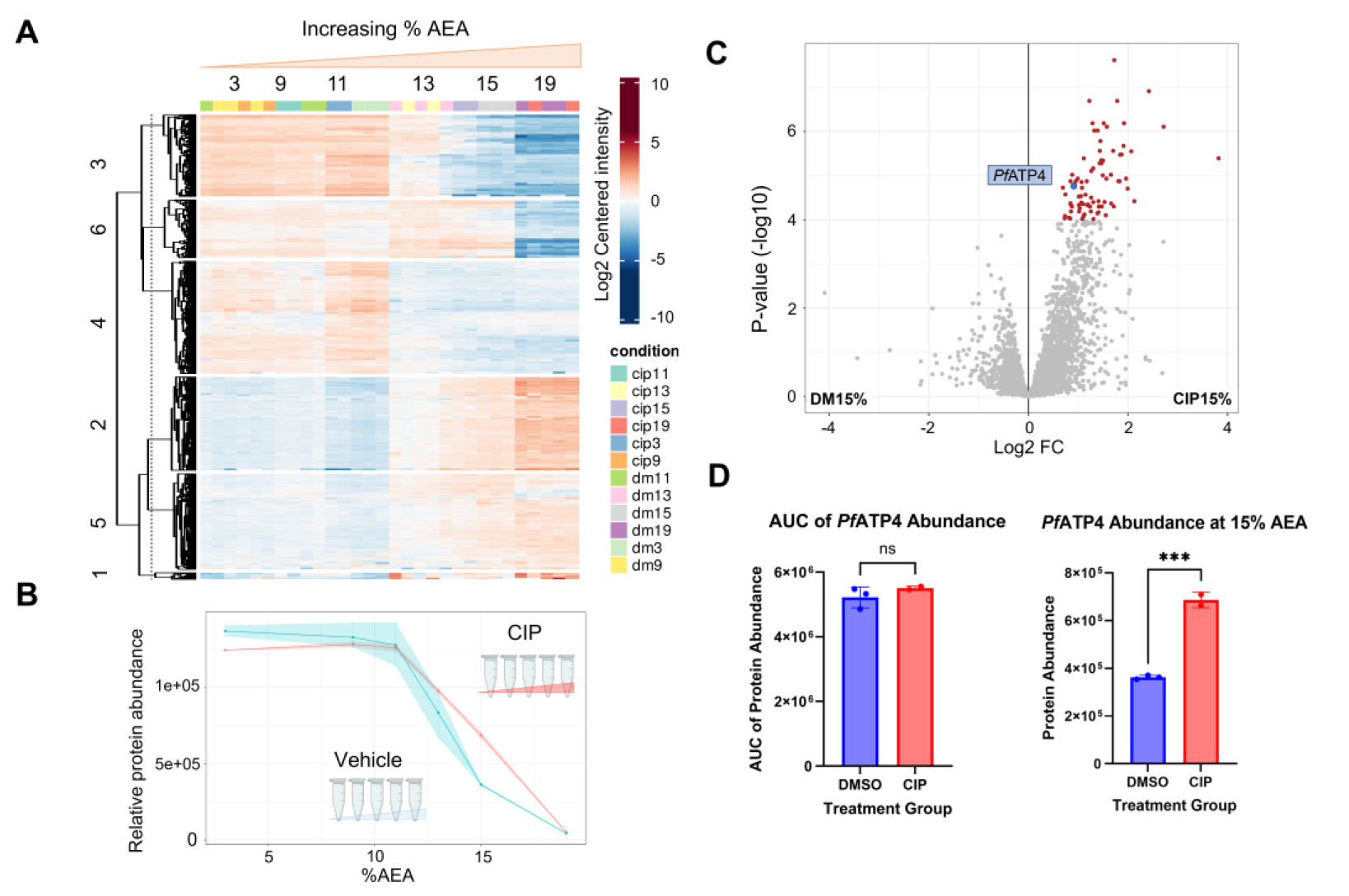
SPP validates protein target of CIP in parasite lysates. (A) Hierarchical clustering of protein abundance changes in parasite lysates treated with increasing concentrations of AEA (3%, 9%, 11%, 13%, 15%, 19%) in the presence of DMSO (vehicle control) or CIP (1 µM). (B) Destabilisation curves of *Pf*ATP4 (PF3D7_1211900) with increasing %AEA, after CIP and DMSO treatments. Protein abundances were plotted against AEA concentrations at 3%, 9%, 11%, 13%, 15%, 19%. The shaded areas around each curve represent the range between the 2-3 replicates at each AEA concentration. (C) Volcano plot comparing vehicle and CIP-treated samples at 15% AEA (vehicle vs drug). *Pf*ATP4, the protein target of CIP, was identified with a log₂FC of 0.92 and *p.adj* of 1.6e-05. Significant proteins shown in red (increased) (*p.adj* < 1e-04, log₂FC > 0.5). (D) Bar plots showing *Pf*ATP4 abundance following DMSO and CIP treatments, based on AUC values and measurements at 15% AEA. The plots were created using GraphPad Prism 10, with error bars representing the standard deviation.

### 3.6 Comparative analysis of multi-drug one-pot SPP in *P. falciparum* lysates

Guided by the CIP results, which underscored the need to analyse solvent percentages individually rather than relying on pooled AUC, we next adopted a “one-pot” strategy to simplify experimental conditions while including positive-control compounds that verify SPP’s performance. One-pot SPP combines multiple drugs in a single lysate and examines their protein target/s in a single SPP assay, for this we used: ATV (targets the cytochrome bc_1_ Complex III), CIP (targets *Pf*ATP4), PYR (targets *Pf*DHFR), MMV1557817 and the synthetic Asn-OSM-S-106 adduct. MMV1557817 has been shown to target *P. falciparum* aminopeptidases M1 (*Pf*A-M1, PF3D7_1311800) and M17 (*Pf*A-M17, PF3D7_1446200) (Vinh et al., 2019), while OSM-S-106 targets *P. falciparum* asparaginyl-tRNA synthetases (*Pf*AsnRS) after forming an enzyme-mediated Asn-OSM-S-106 adduct (Xie et al., 2024), and for both drugs, their targets were previously validated in parasites using various approaches (Edgar et al., 2024; Xie et al., 2024). As enzymatic activation of OSM-S-106 may be impaired in lysate conditions, we used the synthetic Asn-OSM-S-106 conjugate to bypass the need for tRNA-dependent activation (Xie et al., 2024). Most primary targets of the compounds used, including *Pf*DHFR (Figure 5A), *Pf*A-M1, *Pf*A-M17 (Figure 5B), *Pf*ATP4 (Figure 5C), cytoplasmic *Pf*AsnRS (PF3D7_0211800) (Figure 5D) and subunits of the cytochrome bc_1_ Complex III (Figure 5E), exhibited significant stabilisation/destabilisation (*p* < 0.05, |log_2_FC| > 0.5) at different solvent percentages following incubation with the drug mixture in parasite lysate (Table S1). The only exception was CYTC1 (component of cytochrome bc_1_ Complex III targeted by ATV), which showed stabilisation at 5-13 %AEA but did not reach the significance threshold of *p* < 0.05 (at 10% AEA, *p* = 0.30, log₂FC = 0.35) (Table S1, Figure 5E). One-pot SPP validated the targets of earlier single-drug experiments (Figure 2, Figure 3, Figure 4), demonstrating its ability to detect protein targets of multiple drugs at the same time. Moreover, it helps refine target specificity by filtering out off-targets that do not reproducibly stabilise across both single-drug and one-pot SPP. For instance, only MSP1 and ROP were consistently stabilised by CIP (Figure S2), while PYR and ATV yielded no consistently stabilised proteins. The different solvent concentrations at which stabilisation or destabilisation occurs for each protein also highlight the importance of analysing full stability curves across all %AEA to accurately identify drugprotein interactions (Table S1).

**Figure 5.**
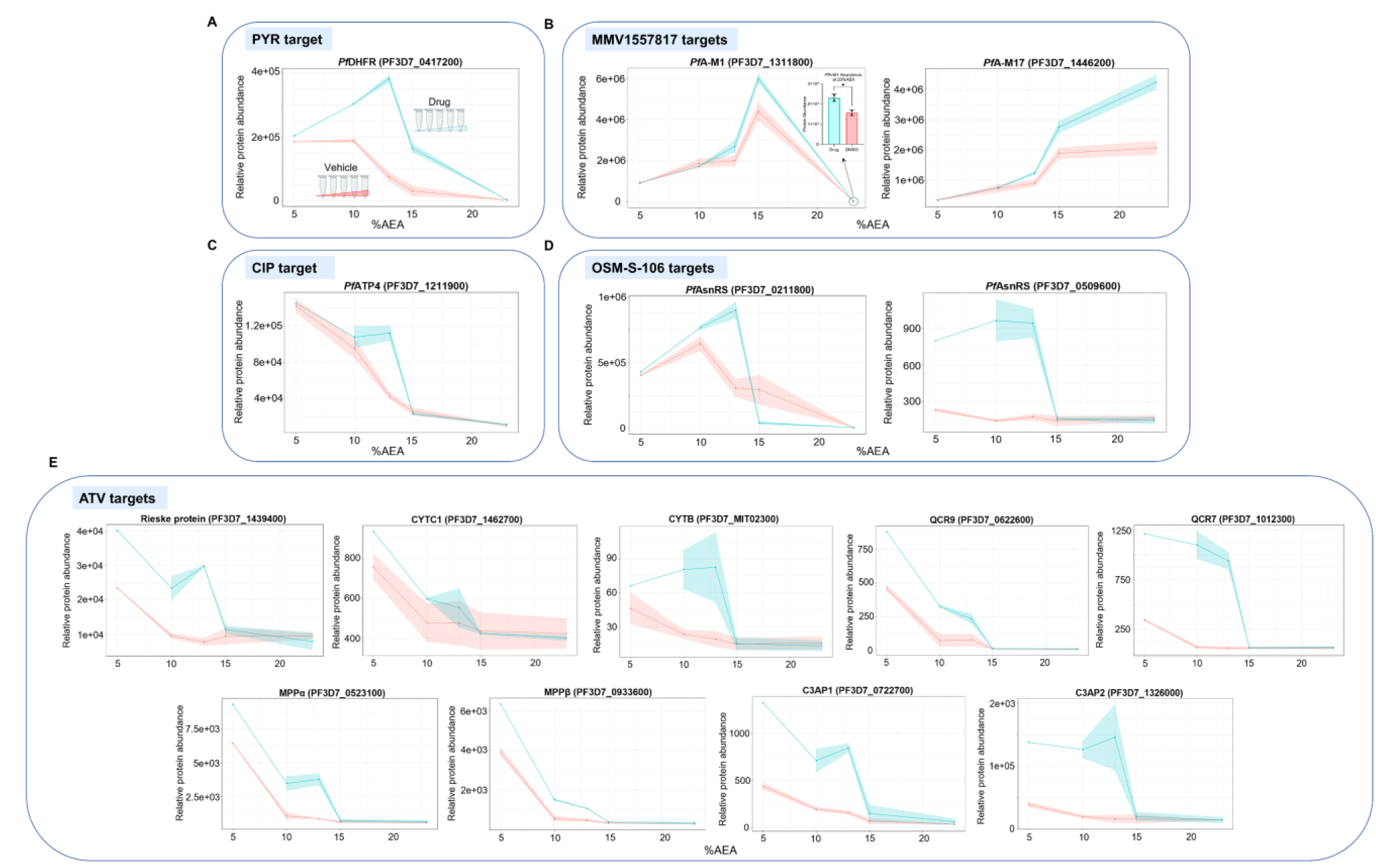
Protein stabilisation profiles from one-pot mixed-drug experiments in *P. falciparum* 3D7 parasite lysates. Curve plots highlighting drug-specific protein targets after drug mixture (1 µM, containing PYR, ATV, CIP, MMV1557817 and Asn-OSM-S-106) and DMSO treatments. Protein abundances were plotted against AEA concentrations (5%, 10%, 13%, 15%, 23%) for key drug targets: (A) *Pf*DHFR (target of PYR). (B) *Pf*A-M1 (PF3D7_1311800) and *Pf*A-M17 (PF3D7_1446200) (targets of MMV1557817). (C) *Pf*ATP4 (target of CIP). (D) Two distinct *Pf*AsnRS (PF3D7_0509600, PF3D7_0211800) (targets of OSM-S-106). (E) Cytochrome bc_1_ Complex III subunits: Rieske protein, CYTC1; CYTB; QCR9; QCR7; MPPα; MPPβ; C3AP1 and C3AP2 (targets of ATV). The shaded areas around each curve represent the range between the 2-3 replicates at each AEA concentration.

*Pf*A-M1 and *Pf*A-M17, two exo-peptidases involved in the final stages of intra-erythrocytic haemoglobin digestion (Goldberg et al., 1990), exhibited modest stabilisation patterns following incubation with the drug mixture containing MMV1557817 (Figure 5B). The one-pot approach applied in our experiments demonstrated stabilisation of *Pf*A-M1 (*p* = 3.12e-02, log₂FC = 0.56) and *Pf*A-M17 (*p* = 1.13e-03, log₂FC = 1.04) by MMV1557817 at 23% AEA, consistent with the known dual-targeting nature of this compound (Edgar et al., 2024). Unlike the *Pf*A-M1 curve which showed destabilisation with higher solvent concentrations, *Pf*A-M17 displayed an opposite trend, where it stabilised as solvent concentration increased (Figure 5B). This difference in stabilisation profile following exposure to AEA could be due to the distinct structures of the two proteins (McGowan et al., 2010; McGowan et al., 2009). Cytoplasmic *Pf*AsnRS exhibited significant stabilisation at 13% AEA (*p* = 1.53e-03, log₂FC = 1.57), but destabilisation at 15% AEA (*p* = 1.28e-05, log₂FC = -2.72) following incubation with the drug mixture containing Asn-OSM-S-106 (Figure 5D), while all other drug targets examined in this experiment showed stabilisation (log_2_FC > 0) (Figure 5). We also identified the apicoplastic *Pf*AsnRS as being stabilised by Asn-OSM-S-106, which was most significant at 13% AEA (*p* = 3.80e-06, log₂FC = 2.75) and showed clear stabilisation only at lower solvent concentrations (5-13% AEA) (Figure 5D). Together, these findings confirm MMV1557817 and OSM-S-106 as inhibitors of *Pf*A-M1, *Pf*A-M17 and two isoforms of *Pf*AsnRS, respectively (Edgar et al., 2024; Xie et al., 2024), supporting their potential as promising antimalarial compounds.

### 3.7 PYR-induced protein target stabilisation following live-cell treatment in *P. falciparum* parasites

Our SPP approach was applied to drug treatments in live parasites, for the first time, and PYR was used as a proof-of-concept compound to assess the differences between live-cell treatment and lysate SPP approaches. Key steps include magnetic enrichment of iRBCs, drug or vehicle treatment, cell lysis and haemoglobin removal using Talon® resin, incubation with increasing %AEA, and analysis via Orbitrap Astral MS using LFQ-DIA proteomics (Figure 6A). The hierarchical clustering heatmap demonstrated the destabilisation of protein groups with increasing AEA, highlighting that AEA induces concentration-dependent effects on the *P. falciparum* proteome obtained from the live-cell treatment SPP approach (Figure 6B). The curve plot for *Pf*DHFR confirmed its stabilisation across AEA percentages following live-cell incubation with PYR (AUC *p.adj* = 2.89e-10, log_2_FC = 0.48) (Figure 6C). A consistent upward shift in the *Pf*DHFR abundance curve was observed across two experiments with live-cell drug treatment (Figure 6, Figure S3), with *Pf*DHFR being the only protein significantly stabilised in both. These results confirm that live-cell treatment SPP is able to identify drug-induced changes in protein stability and can be used for protein target deconvolution. While the stabilisation of *Pf*DHFR was observed following PYR-treatment for both live-cell and lysate experiments, the stability curves of *Pf*DHFR in lysate and live-cell conditions were different (Figure 2C, Figure 6C). At higher percentages of AEA, *Pf*DHFR abundance in drug-free lysates denatured after peaking at 11% (Figure 2C), while in the live-cell treatment study, it increased after 15% (Figure 6C). In contrast to the similar baseline abundances of *Pf*DHFR observed following vehicle and drug treatments in lysate SPP (Figure 2C), live-cell treatment SPP at 5% AEA showed a marginally higher *Pf*DHFR abundance in the PYR-treated samples than in the DMSO controls (Figure 6C). While small, the difference in baseline *Pf*DHFR abundance between drug and vehicle-treated conditions may reflect direct drug-induced stabilisation, or subtle changes in *Pf*DHFR expression or degradation caused by PYR under live-cell conditions.

**Figure 6.**
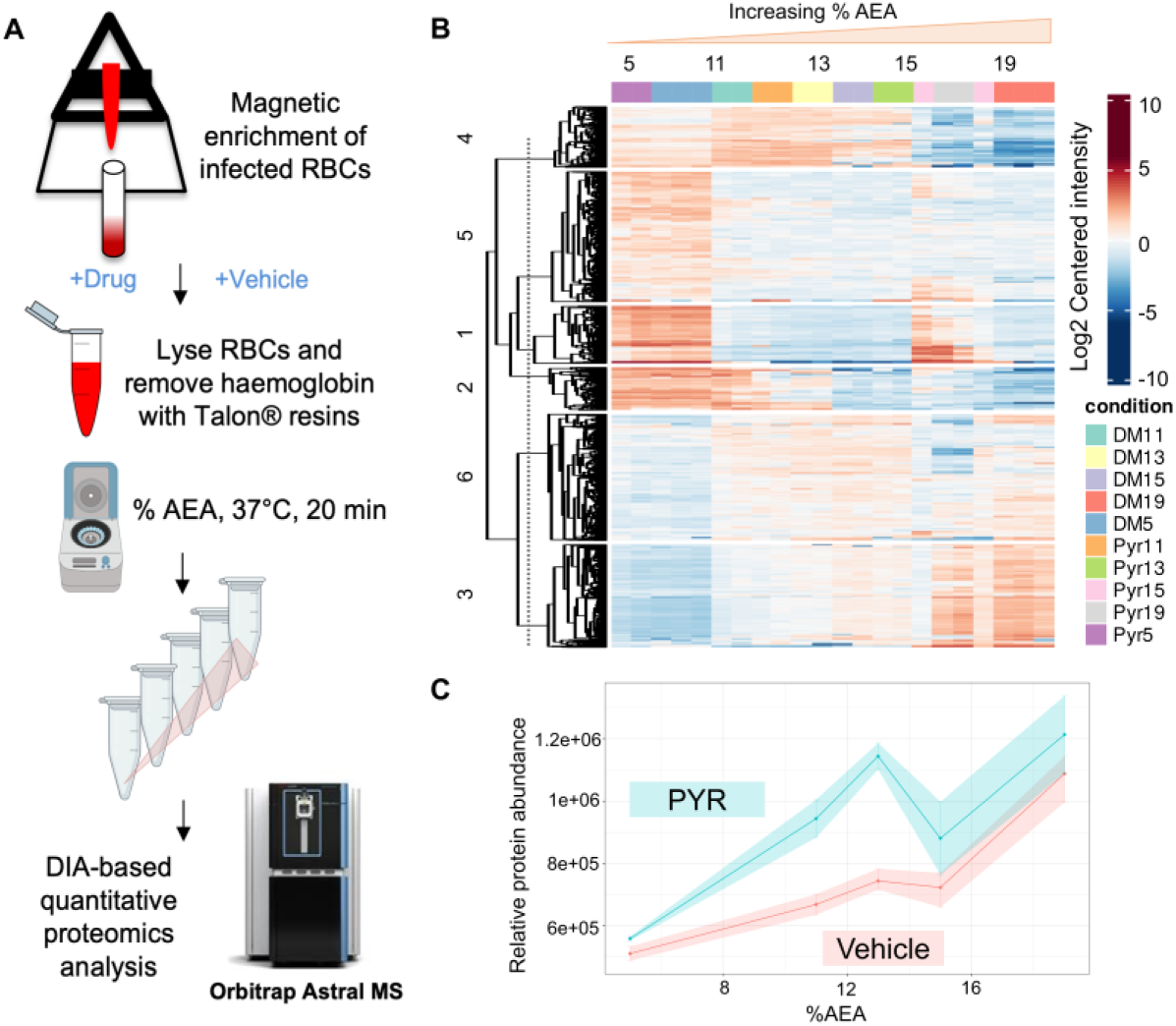
Live-cell treatment SPP of *P. falciparum* 3D7 parasites. (A) Schematic workflow of the live-cell treatment SPP in parasites. Parasites were magnetically enriched and incubated with drug or vehicle, followed by lysis and haemoglobin removal using Talon® resin. Clarified lysates were exposed to increasing %AEA, purified and subjected to downstream proteomic analysis with the Orbitrap Astral MS. (B) Hierarchical clustering heatmap of live parasites treated with increasing concentrations of AEA (3%, 11%, 13%, 15%, 19%) in the presence of DMSO (vehicle control) or PYR (1 µM). (C) Destabilisation curves of *Pf*DHFR with increasing %AEA, after PYR and DMSO treatments. Protein abundances were plotted against AEA concentrations at 3%, 11%, 13%, 15%, 19%. The shaded areas around each curve represent the range between the 2-3 replicates at each AEA concentration.

Since the *Pf*DHFR destabilisation curves differed between lysate and live-cell treatment SPP, we extended our analysis to examine the destabilisation profiles of the remaining proteins in the parasite proteome. A comparative analysis of destabilisation curves of all common parasite proteins identified in both live cell and lysate SPP experiments (2, 771 proteins) (Figure 7A) revealed that 1, 484 proteins, such as transport protein SEC20 (PF3D7_1316800, correlation = 0.998) (Figure 7B), had highly similar or identical stability curves (correlation ≥ 0.75) between lysate and live-cell conditions, accounting for over 50% of the common parasite proteins. The consistent denaturation patterns of closely correlated proteins suggest that SPP can reliably capture protein stability changes for both live cell and lysate incubations. However, a few exceptions exist, where 21.8% of proteins display divergent destabilisation behaviours between the two conditions (correlation < 0). For instance, proliferating cell nuclear antigen 1 (PCNA1, PF3D7_1361900, correlation = -0.749) falls into the negative correlation region (Figure 7C), suggesting that its stability is affected differently in lysates compared to live cell preparations, likely due to differences in protein interactions under the different extraction conditions (George et al., 2024). Similarly, acyl-CoA synthetase (ACS12, PF3D7_0619500) shows moderate correlation (correlation = 0.749) (Figure 7D), suggesting partial differences in its stability profile across the lysate and live-cell treatment SPP protocols, despite similar destabilising trend with increasing %AEA.

**Figure 7.**
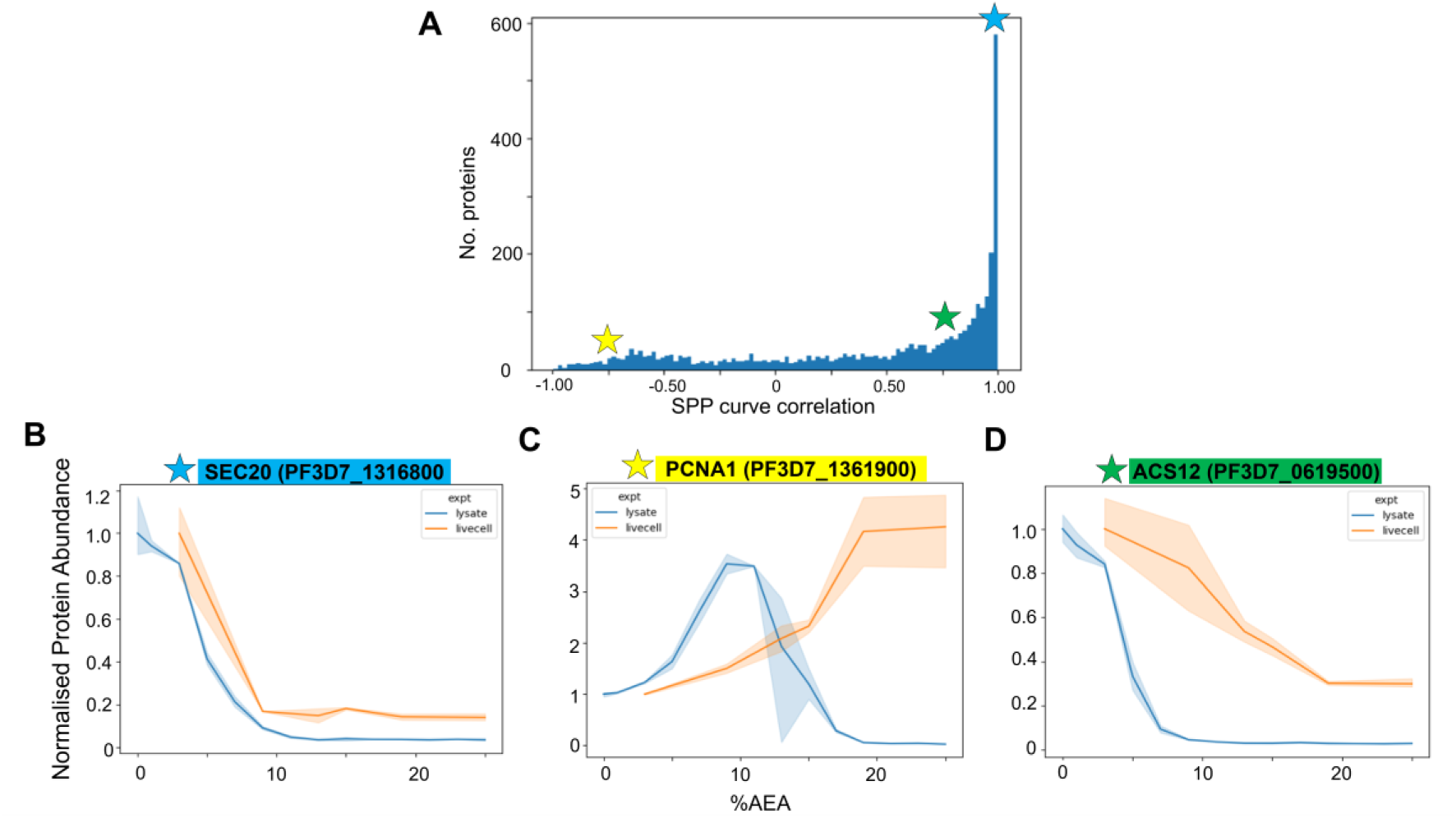
Comparison of destabilisation curves of commonly identified parasite proteins in live-cell treatment and lysate SPP. (A) Distribution of 2, 771 commonly identified parasite proteins in the parasite proteome based on SPP curve correlation between live cell and lysate methodologies. Correlation values range from -1 to 1. Coloured stars mark representative proteins from regions with varying degrees of correlation. (B-D) Representative parasite proteins from distinct correlation regions, protein abundance was normalised against protein abundance in lysates at 0% AEA: (B) Transport protein SEC20 (PF3D7_1316800; correlation = 0.998), representing proteins with highly similar stability profiles across live-cell treatment and lysate SPP (correlation ≥ 0.75). (C) Proliferating cell nuclear antigen 1 (PCNA1, PF3D7_1361900; correlation = -0.749), representing proteins with negative correlation; (D) Acyl-CoA synthetase (ACS12, PF3D7_0619500; correlation = 0.749), representing proteins with low to moderate correlation (0 < correlation < 0.75)

## 4. Discussion

Malaria remains a major global health challenge (WHO, 2014; WHO, 2024), in part due to the rise of drug-resistant parasites (Amaratunga et al., 2016; Balikagala et al., 2021; Gregson and Plowe, 2005; Mairet-Khedim et al., 2021; Phyo et al., 2016; Tumwebaze et al., 2022; Wellems and Plowe, 2001). While many antimalarials exhibit potent activity in phenotypic parasite viability assays, the absence of defined molecular targets limits our ability to predict and mitigate resistance (Challis et al., 2022). Therefore, developing and validating drug target-deconvolution strategies is crucial for combating resistance of existing antimalarials, and supporting the discovery of novel compounds with new targets. In this study, we validated SPP in *P. falciparum* 3D7 lysates as an efficient drug target deconvolution method using a diverse panel of antimalarials, including both approved and preclinical compounds. SPP detects changes in protein stability following drug incubations, which is induced by organic solvents, and this subsequently allows the identification of protein targets. By analysing stability shifts between vehicle- and drug-treated parasite lysates, we identified the respective protein targets of antimalarials with well-characterised targets (PYR, ATV and CIP) and relatively recently discovered antimalarial compounds (MMV1557817 (Vinh et al., 2019) and OSM-S-106 (Gamo et al., 2010)) with high specificity. We further introduced one-pot SPP and live-cell treatment SPP as novel extensions to broaden the applications of the method. One-pot SPP simplifies experimental conditions and provides an extra layer of confirmation by incorporating positive controls, while live-cell treatment SPP has been validated against PYR and captures ligand-protein interactions in a more physiologically relevant context.

The SPP approach builds on recent advances in stability proteomics, such as iSPP, which was recently applied to *P. falciparum* NF54 parasites, and analysed drug-induced stabilisation/destabilisation across asynchronous cultures (Bravo et al., 2024). We proposed and validated live-cell treatment SPP as a novel strategy to capture ligand-protein interactions within intact, infected RBCs. A direct comparison between live-cell treatment and lysate SPP was conducted using PYR, which had previously been examined in Bravo et al. (2024)’s lysate iSPP, making it an ideal candidate for such analysis. Results confirmed that *Pf*DHFR, the primary target of PYR, was consistently stabilised in both live-cell and lysate systems (Figure 2, Figure 6). Notably, in the experiments with livecell PYR incubation, *Pf*DHFR emerged as the sole stabilised target. This level of specificity shows the potential of live-cell treatment SPP as a powerful tool for accurate target ID. One difference between lysate and live-cell treatment SPP is the potential for baseline abundance differences between vehicle and drug-treated samples in live parasites, as observed with *Pf*DHFR abundance at 5% AEA (Figure 6C). This may indicate drug-induced changes in protein expression or turnover during live-cell incubation, and is not entirely reflective of direct ligand-protein binding. To isolate true stability changes, a solvent-free condition (0% AEA) can be incorporated, with all values normalised to the DMSO-treated sample at 0% AEA. This ensures any pre-existing differences before applying AEA are accounted for and only solvent-dependent stabilisation or destabilisation is measured. This also highlights the importance of preserving individual solvent concentrations and generating full destabilisation curves, as such baseline differences would be overlooked if all solvent conditions were combined, as done in iSPP. From a more wholistic perspective, the strong correlation observed between the stability profiles of parasite proteins in live cell and lysate preparations suggests that this approach is likely to be broadly applicable to other druggable protein targets (Figure 7A).

The live-cell treatment approach was proposed to address some limitations of lysate-based SPP. While lysate-based SPP offers a system that is close to the native state, it may not fully capture the complexity of protein interactions in their true cellular environment. Proteins located in certain organelles, such as mitochondria or membrane-bound compartments, can be challenging to study (Evers et al., 2021; Nair et al., 2023). Furthermore, live-cell treatment SPP is ideal for compounds or prodrugs that require intracellular activation before binding to their target. In essence, as an addition to the target deconvolution toolbox, live-cell treatment SPP can provide a more accurate representation of drug-protein interactions within intact cellular contexts.

One-pot mixed-drug SPP is another novelty in our study, where multiple compounds were combined, incubated with one single parasite lysate and tested in one single SPP experiment. This approach significantly reduces sample number while still enabling the detection of multiple ligand-protein interactions within a complex proteome, as evident from the identification of *Pf*DHFR, most subunits of the cytochrome bc_1_ Complex III, *Pf*ATP4, *Pf*A-M1, *Pf*A-M17 and *Pf*AsnRS as respective targets of PYR, ATV, CIP, MMV1557817 and OSM-S-106 (Figure 5). One-pot SPP not only validated the single-drug experiment results of PYR, ATV and CIP (Figure 2, Figure 3, Figure 4), it also identified apicoplastic *Pf*AsnRS as an OSM-S-106 target, which has not been previously reported. A clear caveat of the one-pot approach for target deconvolution of novel compounds is that the stabilised targets cannot be directly linked to individual test compounds. Nevertheless, this approach provides an efficient means to validate initial targets identified in single-drug studies, confirm binding to suspected targets, provide an initial screen of target engagement, or to include drugs with known targets as positive controls alongside novel test compounds for assay quality assurance. Together, these findings substantiate one-pot SPP as a practical and useful approach to improve the efficiency of drug target deconvolution studies.

The combined use of single-drug and one-pot SPP helped eliminate proteins that were not consistently stabilised, thus reducing false discoveries. No off-target proteins were consistently stabilised across both studies for PYR or ATV, whereas only MSP1 and ROP were stabilised following lysate incubation with CIP. However, their localisation and expression are inconsistent with the potent activity of CIP across the parasite intra-erythrocytic lifecycle (Bouwman et al., 2020). MSP1 is predominantly expressed and processed on the surface of extracellular merozoites (Malhotra et al., 2008), and ROPs are synthesised midway through the erythrocytic cycle, but only localised to the rhoptries in merozoites (Sam-Yellowe et al., 1988). Therefore, MSP1 and ROP that are essential for egress and invasion are unlikely to be relevant targets involved in the MOA of CIP (Malhotra et al., 2008; Sam-Yellowe et al., 1988).

The solvent gradient strategy used in these studies was carefully selected based on the denaturation curves observed in our initial profiling (Figure 1), which showed that the majority of *P. falciparum* proteins were mostly denatured by 10% AEA. Therefore, we focused our investigation of drug-induced protein stabilisation on the 9-15% AEA range, compared to the previously established 14-28% AEA (Bravo et al., 2024). While prior iSPP in parasites (Bravo et al., 2024) and human cells (Van Vranken et al., 2021) pooled soluble fractions across solvent conditions to average stability data, our current SPP study individually analysed each solvent percentage. This provides higher resolution protein stability information, as certain ligand-induced stabilisation only occurs at certain chemical environments. For example, *Pf*ATP4 stabilisation by CIP was uniquely observed at 15% AEA (Figure 4). Analysis of individual solvent percentages allows for greater sensitivity and the identification of protein targets that would otherwise be concealed by combining data. This finding suggests that it is particularly critical to take a closer, more detailed look at individual solvent percentages when studying other antimalarials with unknown or poorly characterised targets, where the behaviour of the drug-protein interaction is unpredictable. This unpredictability is also evident where some protein groups increase in abundance with higher AEA concentrations (Figure 1D), with *Pf*A-M17 being an example (Figure 5B). Despite the SPP curve showing increased abundance of *Pf*A-M17 at higher AEA concentrations, a clear difference in soluble protein was still observed between the drug-treated samples and the vehicle control at the higher solvent concentrations.

This study leverages the Orbitrap Astral MS to achieve a coverage of over 3, 800 parasite proteins, accounting for ∼72% of the theoretical 3D7 *P. falciparum* proteome (UniProt ID: UP000001450). To our knowledge, this represents the most extensive malaria proteome coverage reported to date. The Orbitrap Astral MS’s high-resolution accurate mass (HRAM) full scans enable high-speed, high-sensitivity quantification of proteins, including low abundance proteins (Heil et al., 2023; Stewart et al., 2023). This significant increase in coverage did not compromise the quality and precision of data. In contrast to previous single-drug SPP studies, which reported approximately 40 protein hits per experiment (Bravo et al., 2024), our results showed few off-target hits, as no consistently stabilised off-targets were observed for ATV and PYR, while CIP yielded only one or two additional proteins (Figure S2). The results of the SPP study highlight the power and precision of the Orbitrap Astral MS, representing a significant advancement in malaria proteomics.

The results of the SPP approach also align with other protein target identification strategies. The established crystal structures of the cytochrome bc_1_ complex in yeast and *Toxoplasma gondii* demonstrate that ATV engages the catalytic core of Complex III (primarily the Rieske protein, CYTB and CYTC1) (Birth et al., 2014; MacLean et al., 2025). The four accessory proteins of Complex III have been identified in homology searches of apicomplexans (Danne et al., 2012; Hayward and van Dooren, 2019). Direct complexome profiling by Evers et al. (2021) then validated these assignments and established the baseline composition of the complex in *P. falciparum* asexual and sexual stages. There are nine stabilised protein targets of ATV observed using our SPP approach, all of which overlap with previous complexome profiling, including the recently-described apicomplexan-specific components C3AP1 and C3AP2 (Figure 3) (Evers et al., 2021; Maclean et al., 2021). Notably, three additional proteins expected to be part of Complex III showed no ATV-dependent stabilisation: QCR6 was not detected in either our SPP dataset or the complexome profiling, C3AP3 appeared in the Evers et al. (2021) complexome but was absent from our SPP data, and QCR8 was detected in both datasets yet exhibited no significant stability change upon ATV treatment (Figure S1). Variations in protein structure, size, or localisation within Complex III, or experimental procedures could account for the difference in detection or stabilisation of Complex III components (Amporndanai et al., 2018; Birth et al., 2014; Iwata et al., 1998; MacLean et al., 2025; Smith et al., 2012). Whilst the stabilisation of Complex III by ATV does not pinpoint the precise binding site of ATV, this enhanced confidence in the identification of Complex III as the ATV target provides a solid basis for closer investigation of the binding of this test compound to this complex (which for ATV is already well-characterised).

The findings presented in this study demonstrate the utility of SPP profiling as a robust tool for identifying drug targets and understanding drug-protein interactions. However, it is not a one-size-fits-all solution. In biological systems such as human cells, yeast, bacteria, or parasites, their unique structures, metabolic pathways, and proteome complexities give rise to challenges that can lead to variability in target ID through SPP, as seen in previous proteome profiling studies (Bravo et al., 2024; Gygi et al., 2024; Mateus et al., 2018; Van Vranken et al., 2021; Zhang et al., 2020). Additionally, some drugs may not trigger conformational changes large enough to significantly alter protein stability (Gutteridge and Thornton, 2005; Shan et al., 2009; Zheng et al., 2017). SPP, like all target deconvolution approaches, benefits from orthogonal validation, as no single assay captures every interaction. Nevertheless, the optimised SPP workflows described here, with their proteome-wide scope, unbiased target discovery, and diverse applications, represents a valuable addition to the drug discovery toolkit in parasite biology.

## 5. Conclusions

In this study, SPP was successfully applied for target identification in both *P. falciparum* lysates and live cells. Through proof-of-concept experiments using five antimalarials: PYR, ATV, ATV, MMV1557817 and OSM-S-106, we demonstrated the utility of SPP in detecting compound-target engagement. Notably, the novel live-cell treatment SPP was shown to have superior specificity, with the primary drug target (*Pf*DHFR) of PYR precisely identified, highlighting its potential for accurate target deconvolution in a physiologically relevant context. Analysing solvent stability shifts within a refined solvent gradient revealed that certain shifts could only be captured by examining individual solvent percentages. Moreover, the approach enabled detection of the stabilisation of an entire protein complex, providing enhanced confidence in the identification of compound-target interactions. One-pot SPP is a valuable addition by evaluating multiple drugs in one single experimental setup, ideal for secondary validation studies. Looking forward, the integration of single-drug, one-pot and live-cell treatment SPP approaches can be expanded to emerging antimalarials to uncover their novel targets and facilitate the discovery of new drugs for malaria.

## Supplementary Data

Figure S1 shows destabilisation curve plot of QCR8, a previously identified subunit of the cytochrome bc_1_ Complex III targeted by ATV; Figure S2 shows additional proteins stabilised by single-drug treatments with CIP; Figure S3 shows hierarchical clustering and destabilisation curve plot of *Pf*DHFR from a repeat experiment following live-cell PYR treatment; Table S1 lists the protein targets identified in the one-pot mixed-drug SPP. (PDF)

**Supplementary File 1:** Raw proteomics data for solvent-induced protein precipitation; target identification in single-drug SPP experiments with PYR, ATV, and CIP; one-pot lysate SPP with mixed compounds (PYR, ATV, CIP, MMV1557817, and Asn-OSM-S-106); and live-cell treatment SPP experiment with PYR.

## CRediT authorship contribution statement

**Y.J.** and **J.P.M.** contributed equally to this work.

**Y.J.**: investigation, formal analysis, writing – original draft and visualisation, review and editing. **J.P.M.**: investigation, methodology, formal analysis, validation, writing – original draft and visualisation, review and editing. **C.A.M.**: methodology, formal analysis, and writing – review and editing. **H.Z.**: methodology, software, writing – review and editing, data curation. **C.G.**: methodology, and writing – review and editing, **R.B.S.**: methodology, software, data curation, writing – review and editing, supervision. **D.J.C.**: methodology, formal analysis, and writing – review and editing, supervision, project administration, and funding acquisition. **G.S.**: investigation, methodology, formal analysis, conceptualisation, writing – review and editing, supervision, project administration, and funding acquisition.

## Notes

The authors declare no competing financial interest.

## Acknowledgment

We gratefully acknowledge funding support from the Medical Research Future Fund (Drug Target Identification Platform; MRFCRI000102) and ARC Future Fellowship to DJC (FT220100564). Analysis was performed in the Drug Target Identification node of the Monash Proteomics and Metabolomics Platform. This study was further supported by Monash eResearch capabilities, including Research Data Storage and Nectar Research Cloud.

## Abbreviations

SPP: solvent proteome profiling
iSPP: integral solvent-induced proteome profiling
LC-MS/MS: liquid chromatography-mass spectrometry/mass spectrometry
*Pf*: Plasmodium falciparum
Target ID: target identification
ACTs: artemisinin-based combination therapies
ART: artemisinin
RBC: red blood cell
iRBC: infected red blood cell
SPROX: stability of proteins from rates of oxidation
AEA: acetone/ethanol/acetic acid (50:50:0.1 v/v/v)
AUC: area under the curve
BH: Benjamini-Hochberg
FC: fold change
MSP: merozoite surface protein
ROP: putative rhoptry protein
ATP4: ATPase 4
tRNA: transfer ribonucleic acid
QCR6: cytochrome bc_1_ complex putative subunit 7
QCR7: cytochrome bc_1_ complex putative subunit 7
QCR8: cytochrome bc_1_ complex putative subunit 8
QCR9: cytochrome bc_1_ complex putative subunit 9
CYTB: cytochrome B
CYTC1: cytochrome c1
C3AP1: respiratory chain complex 3 associated protein 1
C3AP2: respiratory chain complex 3 associated protein 2
C3AP3: respiratory chain complex 3 associated protein 3
MPPα: mitochondrial-processing peptidase subunit alpha
MPPβ: mitochondrial-processing peptidase subunit beta
DIA: data-independent acquisition
LiP-MS: limited proteolysis-coupled mass spectrometry
PPIs: protein-protein interactions
iRT: indexed retention time
FA: formic acid
ACN: acetonitrile
MOA: mechanism of action
DHFR: bifunctional dihydrofolate reductase-thymidylate synthase
A-M1: aminopeptidase M1
A-M17: aminopeptidase M17
AsnRS: cytoplasmic asparaginyl-tRNA synthetase
PCNA1: proliferating cell nuclear antigen 1
ACS12: acyl-CoA synthetase
ATV: atovaquone
CIP: cipargamin
PYR: pyrimethamine
DMSO/DM: dimethyl sulfoxide
TPP: thermal proteome profiling
PCA: principal component analysis
CETSA: cellular thermal shift assay
LFQ-DIA: label-free quantification in data-independent acquisition
MS: mass spectrometry/mass spectrometer
*p.adj*: adjusted *p* value
RSD: relative standard deviation
HRAM: high-resolution accurate mass.

## Notes

### Competing Interest Statement

The authors have declared no competing interest.

