## Supplementary Figures for "Validation of Solvent Proteome Profiling for Antimalarial Drug Target Deconvolution"

### Supplementary Data

**Figure S1**. Abundance curve of previously identified cytochrome $bc_1$ complex III subunit QCR8 (PF3D7_0306000), following atovaquone (ATV) treatment in parasite lysate (single-drug)

**Figure S2.** Destabilisation curves of additional *P. falciparum* proteins stabilised by cipargamin (CIP) treatment in parasite lysate (single-drug)

**Figure S3.** Hierarchical clustering heatmap and *Pf*DHFR abundance curve following PYR incubation in an independent repeat of live-cell treatment SPP

**Table S1.** Protein targets of antimalarials in mixed-drug one-pot SPP

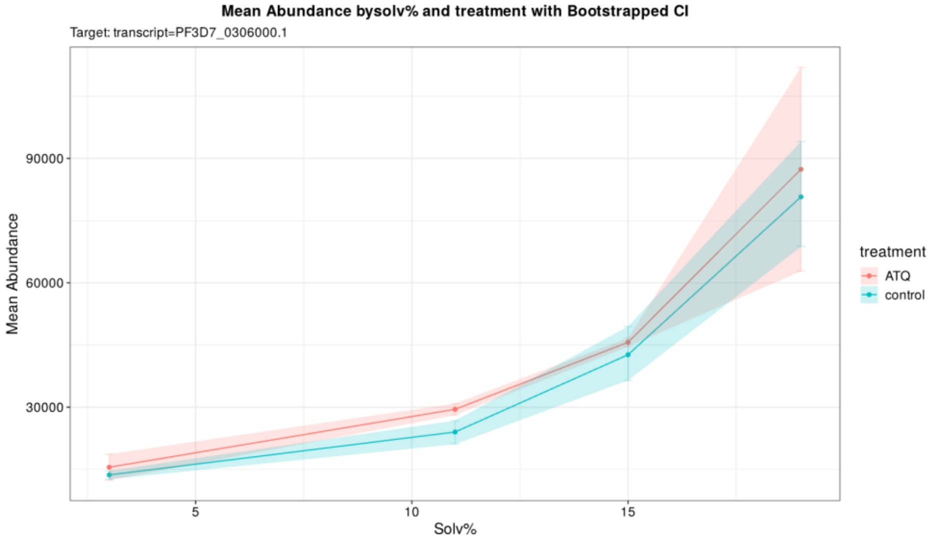


**Figure S1**. **Abundance curve of cytochrome bc$_1$ complex subunit 8 (QCR8, PF3D7_0306000) after ATV (1 µM) treatment in single-drug lysate SPP.** Protein abundances were plotted against AEA concentrations at 3%, 11%, 15%, 19%. AUC analysis revealed non-significant stabilisation (*p.adj* > 0.05). The shaded areas around each curve represent the range between the 2 replicates at each AEA concentration.

A

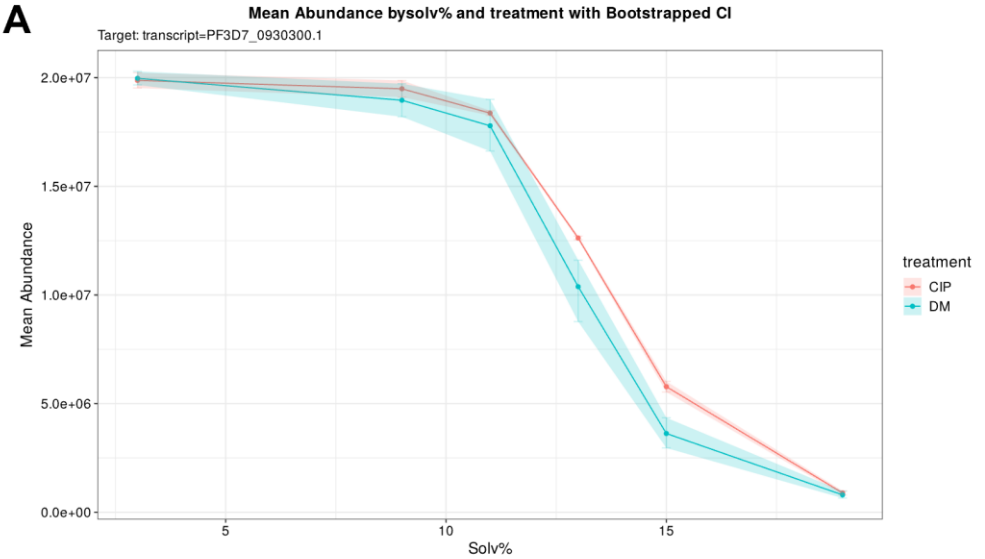


B

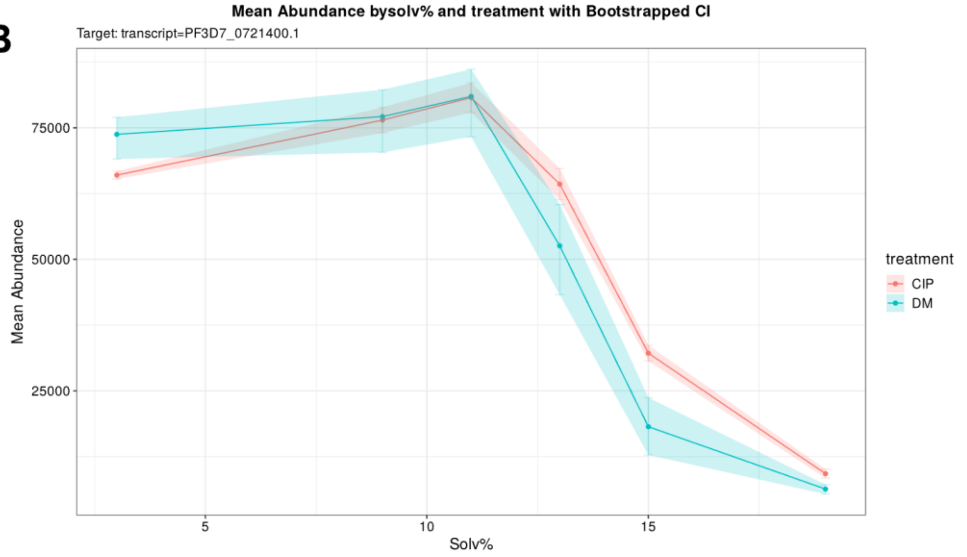


**Figure S2. Destabilisation curves of additional *P. falciparum* proteins that showed stabilisation following CIP treatment in lysate SPP.** Protein abundances were plotted against AEA concentrations at 3%, 11%, 15%, 19%. (A) merozoite surface protein 1 (MSP1, PF3D7_0930300) exhibited significant stabilisation upon incubation with 1 µM CIP in parasite lysates (at 15% AEA, *p.adj* = 1.33e-05; $\log_2$FC = 0.95); (B) putative rhoptry protein (ROP, PF3D7_0721400) exhibited significant stabilisation upon incubation with 1 µM CIP in parasite lysates (at 15% AEA, *p.adj* = 6.85e-06; $\log_2$FC = 1.3). The shaded areas around each curve represent the range between the 2 replicates at each AEA concentration.

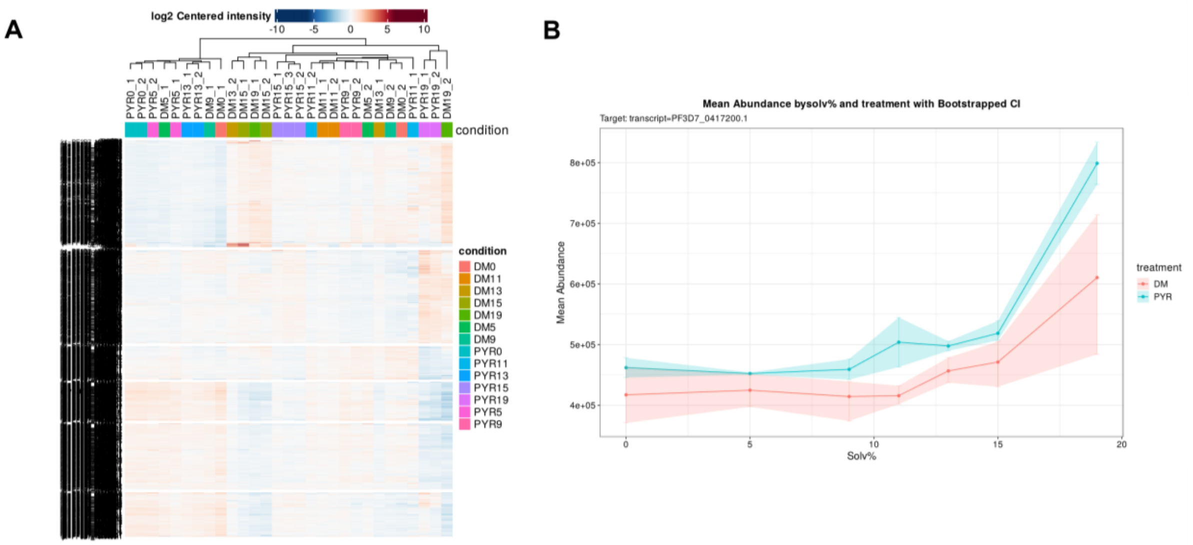


**Figure S3. Live-cell treatment SPP repeat using pyrimethamine (PYR).** (A) Hierarchical clustering analysis illustrating distinct protein group responses to increasing AEA concentrations, highlighting patterns of stabilisation and destabilisation across samples. (B) Destabilisation curve of *Pf*DHFR (PF3D7_0417200), the primary target of PYR, showing AEA concentration-dependent stabilisation following live-cell incubation with PYR (10 µM). Protein abundance was plotted across AEA concentrations at 0%, 5%, 9%, 11%, 13%, 15%, and 19%. The most significant stabilisation was observed at 19% AEA (*p.adj* = 1.61e-02; $log_2$FC = 0.53). The shaded area around each curve represents the range between the two replicates at each AEA concentration.

**Table S1. Proteins identified with a shift in abundance in one-pot SPP experiments across different solvent percentages[a]**

| Antimalarial | Protein target | %AEA with the most significant stability shift | Drug vs DMSO $log_2FC$ at that %AEA | *p*-value |
|---|---|---|---|---|
| **OSM-S-106** | Cytoplasmic *Pf*AsnRS[b] | 15% | -2.72 | 1.28e-05 |
| | Apicoplastic *Pf*AsnRS[c] | 13% | 2.75 | 3.80e-06 |
| **PYR** | *Pf*DHFR[d] | 15% | 2.63 | 3.93e-05 |
| **ATV** | CYTB[e] | 13% | 2.11 | 5.93e-03 |
| | C3AP1[f] | 13% | 2.4 | 2.42e-03 |
| | C3AP2[g] | 13% | 3.21 | 1.07e-04 |
| | MPPα[h] | 13% | 2.12 | 1.67e-05 |
| | MPPβ[i] | 10% | 1.53 | 1.5e-04 |
| | *CYTC1[j] | 10% | 0.35 | 0.30 |
| | QCR9[k] | 10% | 2.51 | 5.01e-03 |
| | QCR7[l] | 10% | 4.05 | 1.67e-07 |
| | Rieske protein[m] | 13% | 1.94 | 9.24e-05 |
| **CIP** | *Pf*ATP4[n] | 13% | 1.4 | 1.04e-04 |
| **MMV1557817** | *Pf*A-M1[o] | 23% | 0.56 | 3.12e-02 |
| | *Pf*A-M17[p] | 23% | 1.04 | 1.13e-03 |

[a] Data are presented as the solvent percentage with the most significant stability shift for each protein in one-pot SPP experiment, along with the corresponding $log_2FC$ and *p*-value. Proteins listed include known drug targets of PYR, ATV, CIP, MMV1557817 and OSM-S-106 (Asn-OSM-S-106 was used in the experiment), both significantly and non-significantly stabilised (which was marked with a *). *p*-values below 0.05 were considered significant.

[b] *Pf*AsnRS (PF3D7_0211800): Asparaginyl-tRNA synthetase (cytoplasmic)

[c] *Pf*AsnRS (PF3D7_0509600): Asparaginyl-tRNA synthetase (apicoplastic)

[d] *Pf*DHFR (PF3D7_0417200): Bifunctional dihydrofolate reductase-thymidylate synthase

[e] CYTB (PF3D7_MIT02300): Cytochrome B

[f] C3AP1 (PF3D7_0722700): Respiratory chain complex 3 associated protein 1

[g] C3AP2 (PF3D7_1326000): Respiratory chain complex 3 associated protein 2

[h] MPPα (PF3D7_0523100): Mitochondrial-processing peptidase subunit alpha

[i] MPPβ (PF3D7_0933600): Mitochondrial-processing peptidase subunit beta

[j] CYTC1 (PF3D7_1462700): Cytochrome $c_1$

[k] QCR9 (PF3D7_0622600): Cytochrome $bc_1$ subunit 9

[l] QCR7 (PF3D7_1012300): Cytochrome $bc_1$ subunit 7

[m] Rieske protein (PF3D7_1439400) - Cytochrome $bc_1$ rieske protein

[n] *Pf*ATP4 (PF3D7_1211900): Non-SERCA-type Ca2+ -transporting P-ATPase

[o] *Pf*A-M1 (PF3D7_1311800): Aminopeptidase M1

[p] *Pf*A-M17 (PF3D7_1446200): Aminopeptidase M17
